# Toward Optimizing Thalamic Deep Brain Stimulation for Cortical Modulation: A Surrogate Brain Approach

**DOI:** 10.64898/2026.06.26.734900

**Authors:** Raunak Ahmed, Yuqi Feng, Anna Wang Roe, Zhe Sage Chen

## Abstract

The thalamus is a central hub that interfaces with widespread cortical and subcortical nodes. Thalamic deep brain stimulation (DBS) offers a principled strategy for distributed cortical modulation: since distinct thalamic nuclei project to spatially segregated cortical territories, stimulation at a single thalamic site can influence multiple cortical nodes. Realizing this potential requires accurate subject-specific estimates of directed thalamocortical effective connectivity (EC) and a computational framework for optimizing stimulation parameters that achieve desired cortical responses. Here, we address both challenges using Neural Perturbational Inference (NPI), a surrogate-brain approach that estimates EC by applying virtual perturbations to a nonlinear dynamical model fitted to resting-state fMRI data. We extend NPI to a high-resolution thalamocortical network comprising 360 cortical regions and 442 thalamic voxels spanning 12 nuclei. We introduce two innovations in training: (i) a temporal signal-to-noise ratio (tSNR)-weighted loss accounting for signal heterogeneity, and (ii) a multi-resolution, cross-scale consistency loss that regularizes model complexity. These strategies yield improved performance in synthetic benchmarks across varying tSNR regimes. Leveraging the inferred subject-specific EC, we further formulate a constrained linear control problem to identify sparse thalamic stimulation targets that achieve desired cortical activation patterns. We validate the inferred EC structure on two independent datasets: the MacStim dataset comprising two macaque monkeys with infrared neural stimulation on medial pulvinar, and the HumanTC resting-state fMRI dataset comprising twelve human subjects. Our results reveal site-specific thalamocortical EC profiles, producing interpretable predictions that align with known ground-truth structures. Together, this work establishes a computationally grounded pathway toward personalized optimization of thalamic DBS in both human and nonhuman primates.

## 1 Introduction

Thalamic deep brain stimulation (DBS) is clinically established for Parkinson’s tremor [1, 2] and disorders of consciousness [3]. A key property of the thalamus is that individual nuclei project to distinct, large cortical territories [4], so stimulating a single thalamic site can simultaneously modulate multiple cortical targets — a distributed control strategy unavailable through direct cortical stimulation. Translating this into principled DBS parameter selection requires a computational strategy that enables directed causal inference in thalamocortical network.

In neuroscience, structural connectivity (SC) maps the physical wiring of the brain and functional connectivity (FC) captures statistical dependencies among neural activities. In contrast, effective connectivity (EC) captures directed, causal dependencies between nodes in the brain network [5–7]: EC_*ij*_ quantifies how much target region *j* responds when source region *i* is perturbed. EC can be derived through neurostimulation experiments [8] or statistical methods (such as dynamical causal modeling and Granger causality), which are often restricted to analysis of small networks [9, 10]. Recently, a surrogate brain approach known as Neural Perturbational Inference (NPI) has been developed for EC inference based on resting-state fMRI-BOLD signals [11]; NPI was validated only on 360 cortical ROIs in synthetic data, but has neither been tested in a large thalamocortical network nor been combined with causal neurostimulation perturbations.

Our paper features several key contributions.

1. **Thalamocortical EC at scale**. We extend NPI to 802 nodes (360 cortical ROIs from MMP atlas plus 442 voxels derived from the thalamus mask from THOMAS segmentation)—the very first large thalamocortical EC map at the voxel-level resolution from resting-state fMRI. We further use the inferred EC to identify the hub in the thalamocortical network.
2. **DBS optimization framework**. The inferred EC serves as an approximate linear influence model for solving a sparse LASSO target-selection problem; the goal of optimization is to find a minimal set of thalamic nuclei and associated stimulation intensity that yields a desired cortical response. We validate the proof-of-concept strategy using the Macaque monkey dataset.
3. **SNR-weighted loss (Propositions 1–2) and multi-resolution training (Theorem 1)**. We propose a principled heteroscedastic noise model that addresses the lower temporal SNR (tSNR) in deep thalamic nuclei. We also develop a cross-scale consistency loss function that reduces effective model complexity. Both theoretical proof and empirical validation are provided.

## 2 Related Work

### Effective connectivity

Resting-state EC has been widely studied in brain networks [12]. In terms of methodology, DCM [9, 13] is not-scalable and limited to a small number of nodes. Granger causality

[10] is slightly more scalable but cannot handle dimensionality at the order of hundreds; it may also conflate statistical with causal dependencies under hidden-state fMRI models [14]. NPI replaces biophysical parameterization with a data-driven approach and was validated in a large cortical network with fMRI data [11].

### Thalamocortical connectivity

Thalamocortical FC has been studied via tractography and resting-state fMRI data [4, 15]. While large-scale functional networks provide the mesoscale context in which thalamic nuclei operate, directed EC at the voxel resolution has not been estimated from neuroimaging data. The THOMAS parcellation [16] provides a reproducible 12-nucleus-per-hemisphere atlas used in our analysis.

### Surrogate brain models

To accommodate a large fine-resolution thalamocortical network (802 nodes), we proposed DropoutMLP regularization [17] with two new loss functions in training. Alternative sequence models and self-supervised representation learning approaches [18, 19] offer richer inductive biases but are not adopted here given a relatively small sample size.

### Computational DBS planning

Atlas-based and diffusion-MRI approaches capture anatomical projections but not directed causal influence [20]. The digital twin brain (DTB) or virtual brain framework uses a similar idea but builds upon biophysical models [21, 22], yet our data-driven approach targets the thalamic stimulation for distributed cortical modulation. Our framework (see Figure 1) offers a closed-form convex optimization for principled stimulation parameter selection.

**Figure 1:**
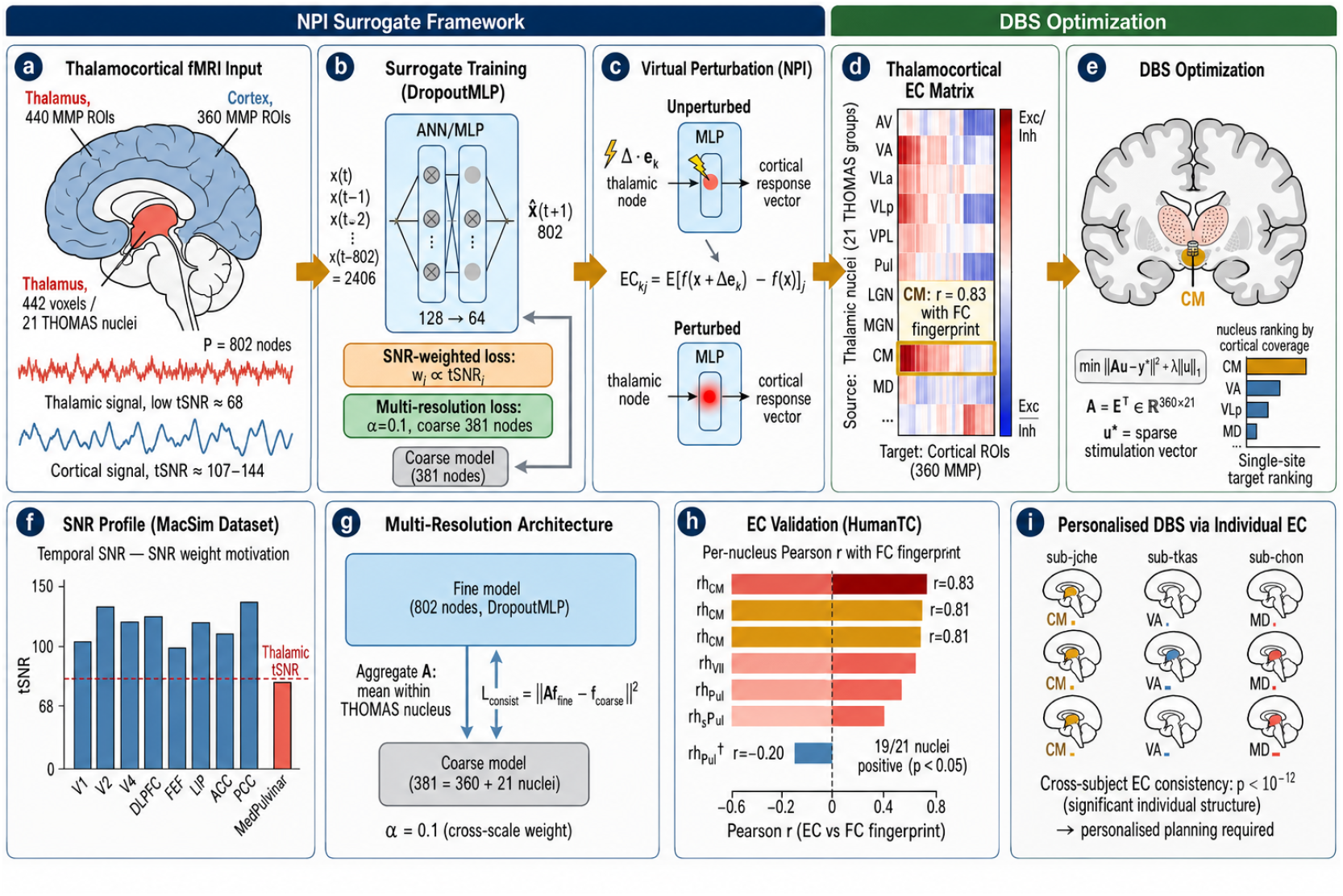
Overview of the proposed framework. **(a)** Large thalamocortical network: 360 cortical MMP ROIs and 442 thalamic voxels (21 THOMAS nuclei). **(b)** Surrogate brain model: a DropoutMLP is trained on sliding-window BOLD sequences with two extensions: an SNR-weighted loss that down-weights low-SNR thalamic voxels, and a multi-resolution consistency loss that jointly trains a fine-resolution (802-node) and coarse-resolution (381-node) model. **(c)** Virtual node perturbation in the surrogate brain model. **(d)** Thalamocortical EC matrix (21 THOMAS nuclei × 360 cortical ROIs); warm/cool color: excitatory/inhibitory. **(e)** DBS optimization. **(f)** Distinct tSNR profile motivating the use of weighted loss. **(g)** Schematic of multi-resolution architecture. **(h)** Per-nucleus EC validation against FC fingerprint (HumanTC fMRI dataset). **(i)** Personalized DBS: individual EC maps vary across subjects, yielding subject-specific stimulation targets.

## 3 Methods

### 3.1 NPI Framework

NPI employs self-supervised learning to train a surrogate artificial neural network (ANN) such as multilayer perceptron (MLP) *f*_*θ*_ : ℝ^*N s*^→ ℝ^*N*^ to predict the next fMRI-BOLD frame from a history of *s* consecutive frames (using mean-squared error), where *N* is the number of brain nodes and *s* = 3 throughout this work. Next, NPI virtually perturbs each node to estimate the EC:

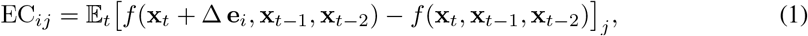

where **e**_*i*_ a unit vector, and Δ = 0.5×std(BOLD). Differences in predicted target region responses, compared to baseline inputs, reflect the EC from the source (perturbed) to the target regions; increased or decreased activity in the target regions indicates excitatory or inhibitory EC, respectively. One-to-all EC mapping is achieved by perturbing a single node, and systematic perturbations across all nodes provide an all-to-all EC mapping.

### 3.2 DropoutMLP Architecture

To improve generalization in training ANNs on real fMRI data, we imposed regularization on the standard MLP network architecture. To mitigate this overfitting problem, we introduced **DropoutMLP** [17] with the following setup: batch size 64, dropout probability *p* = 0.2 after each ReLU layer, Adam optimizer (lr = 3 × 10^*−*4^). To monitor overfitting, we employed early stopping (patience 40) and computed the loss ratio between testing and training sets across all subjects.

### 3.3 SNR-Weighted Training Loss

Since deep thalamic nuclei exhibit lower temporal SNR (tSNR) than the cortex, we modeled fMRI-BOLD observations as

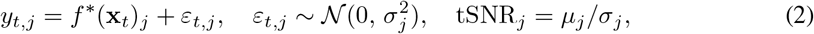

where *µ*_*j*_ denotes the empirical mean BOLD signal of region *j*. The maximum-likelihood loss under this model is a *weighted* MSE:

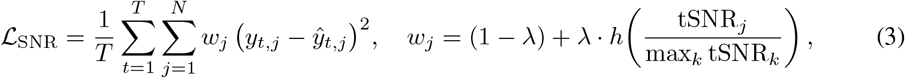

where *h* : [0, 1] → [0, 1] is monotonically non-decreasing, *N* denotes the size of the output, and *λ* ∈ [0, 1] controls the strength of SNR weighting. High-tSNR regions receive weight ≈ 1, whereas low-tSNR regions are down-weighted toward (1 − *λ*). We used *λ* = 0.5 and *h*(*u*) = *u* throughout.

### 3.4 Multi-Resolution Training

We trained an MLP at a fine-resolution voxel level (loss ℒ_fine_) and further augmented it with a cross-scale consistency loss (ℒ_cross_) at the nucleus-coarse resolution:

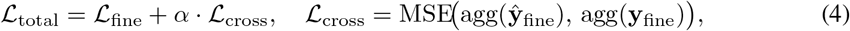

where agg(·) averages voxel predictions within the thalamic nucleus group. The cross-scale loss acts as a structured regularizer, reducing the effective model complexity (Theorem 1). We chose *α* via grid search within a logarithmically spacing range to obtain the best generalization.

### 3.5 Quantitative Assessment

Upon the convergence of ANN training, we assessed the model performance on held-out test data. In the synthetic dataset, Pearson’s correlation was computed between the NPI-inferred EC and the ground-truth EC (GT-EC). In the monkey fMRI dataset, because of the lack of GT-EC, we estimated the correlation between model-predicted cortical response and experimentally perturbed cortical responses through thalamic neurostimulations. In the human fMRI dataset, we computed the correlation between the model-based FC and empirical FC a the voxel level and compared the population-averaged EC with an independent FC fingerprint map. Given the inferred EC graphs from each dataset, we computed graph-theoretic nodal centrality metrics, such as in-degree (influence recipients) and out-degree (influence sources), and identified the “hub” of the network based on the rank of sum of in-degree and out-degree for all nodes.

## 4 Theoretical Analysis

Here we provide theoretical arguments for the SNR-weighted loss (Propositions 1–2) and multi-resolution training (Theorem 1). Full proofs are given in Appendix A.

### Proposition 1

(SNR weighting: unbiasedness and variance reduction). *Modelling BOLD as* 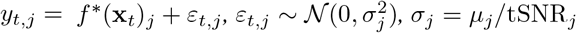, *the SNR-weighted least-squares estimator*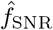 *is unbiased for f*^*∗*^ *and satisfies*

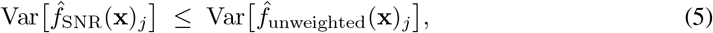

*with the reduction factor equal to* 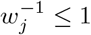, *where w*_*j*_ *is monotonically non-decreasing in* tSNR_*j*_.

### Proposition 2

(Generalization bound under SNR weighting). *With probability* 1− *δ, the excess risk of* 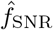 *over the Bayes-optimal f*^*∗*^ *satisfies*

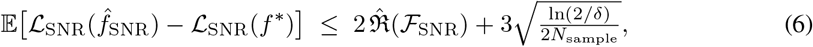

*Where* 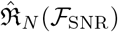 *is the empirical Rademacher complexity of the SNR-weighted function class* F_SNR_ = {(*f*, **x**) → Σ_*j*_ *w*_*j*_(*f* (**x**)_*j*_ − *y*_*j*_)^2^ : *f* ∈ F} *[23]*.

### Theorem 1

(Multi-resolution complexity reduction). *Let K denote the number of thalamic nucleus, and N denotes the total number of cortical and thalamic voxels in the thalamocortical network; let ℱbe a class of MLP output functions f* (**x**) = *W*_out_*ϕ*(**x**) *with trace-norm bound* ∥*W*_out_∥_*∗*_≤ *B. Under a uniform trace-norm distribution across the N output dimensions, the K nucleus-level consistency constraints in Eq. (4) reduce the effective Rademacher complexity as follows:*

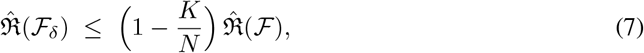

*and the number of samples required to achieve generalization error* ≤ *ε with probability* ≥ 1 − *δ*_0_ *satisfies: N*_multi*−*resolution_ ≤ (1 − *K/N*)^2^ *N*_sample_.

*Remark:* With *K* = 21 THOMAS nuclei and *N* = 802 total nodes in the human fMRI dataset: complexity reduction *K/N* ≈ 0.026, giving *N*_multi*−*resolution_ ≤ 0.949 *N*_sample_.

## 5 Experiments and Results

### 5.1 Synthetic Data Validation

We first validated our proposed method in a fully controlled setting where ground-truth EC is known. By adapting the strategy of [11], we first constructed a 50-node thalamocortical recurrent neural network (RNN) (*N*_thal_ = 15, *N*_cort_ = 35) and generated synthetic data under five progressively challenging SNR scenarios. Next, we varied the number of total nodes (*N* = 20, 50, 100, 200, 300) and simulated sample size (*N*_sample_ = 1, 000, 2, 000, 5, 000, 10, 000, 20, 000) and assessed the model robustness. See Appendix B for model training details. We considered three SNR conditions:

1. **Uniform**: SNR is uniform across all nodes.
2. **Moderate**: thalamic nodes have a mean tSNR of 5.5; additional smoothing (Gaussian variance 1) is applied.
3. **Severe**: thalamic nodes have a mean tSNR of 3.5.

For each SNR condition, we examined three loss functions (standard, weighted SNR, multi-resolution) and compared the Pearson’s correlation between the estimated and ground-truth EC.

In the case of *N* = 50, the proposed weighted SNR function slightly outperformed the standard loss function (Table 1) in the moderate and severe SNR conditions (Fig. 2a). The multi-resolution model was probably overfitting when the training samples were insufficient, yielding slightly poorer generalization; but its performance improved when the sample size increased. Varying the sample size and network dimensionality showed that the results improved with more data (Fig. 2c), supporting the robustness of the method in a controlled setting. Furthermore, in both moderate and severe SNR conditions, the weighted SNR loss achieved the smallest training/testing loss gap, consistent with the theoretical prediction.

**Table 1:** Pearson’s correlation (mean ± s.d. based on 5-fold cross-validation with different random seeds) between the estimated EC and ground truth under three SNR conditions.

| Method | Uniform SNR | Moderate SNR | Severe SNR |
| --- | --- | --- | --- |
| Standard | **0.744±0.007** | 0.827±0.005 | 0.832±0.003 |
| Weighted SNR | 0.738±0.007 | **0.829±0.004** | **0.833±0.004** |
| Multi-Resolution | 0.733±0.003 | 0.825±0.003 | 0.825±0.002 |

**Figure 2:**
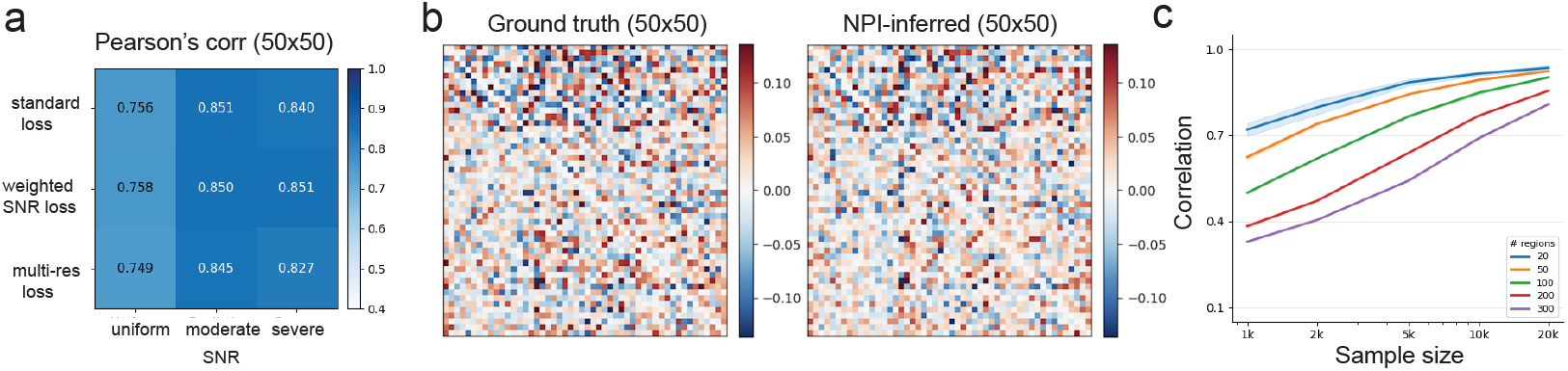
Synthetic data results. (**a**) EC fidelity comparisons under three SNR conditions (result from one fold in cross-validation). (**b**) Ground-truth vs NPI-inferred EC. (**c**) Comparison of Pearson’s correlation with various simulated network sizes and sample sizes. All illustrations were obtained with weighted SNR loss and moderate SNR condition.

### 5.2 Dataset 1: Macaque Monkey fMRI Recordings

#### Data

The Macaque Monkey Thalamic Stimulation dataset (MacStim) consists of two macaque monkeys (P and Y) recorded at four fMRI sessions (7T MAGNETOM scanner; Siemens Healthineers) [24]: (i) a resting-state session (240 time points or TPs, equivalent to ≈ 8 min with 0.5 Hz sampling frequency) used for ANN training, and (ii) three held-out infrared neural stimulation sessions at distinct medial pulvinar (PM) sites — P1 (720 TPs, 45 trials), P2 (480 TPs, 30 trials) and P3 (720 TPs, 45 trials) as well as Y1 (30 trials), Y2 (30 trials) and Y3 (45 trials). Eight cortical regions were included: V1, V2, V4 (visual hierarchy), DLPFC (dorsolateral prefrontal cortex), FEF (frontal eye field), LIP (lateral intraparietal cortex), ACC (anterior cingulate cortex), PCC (posterior cingulate cortex), plus three thalamic sites at the medial pulvinar (PM), yielding a 11-node thalamocortical network. Raw BOLD values were demeaned per region before training. Experimental procedure was approved by the Institutional Animal Care and Use Committee (IACUC) of Zhejiang University [24].

#### Temporal SNR profile

tSNR (defined as mean/std on raw fMRI-BOLD before normalization) was measured per region. All cortical regions showed comparable mean tSNR (e.g., monkey P: V1: 107, V2/LIP: 130, ACC: 144); the SNR of PM is markedly lower (tSNR = 68.23).

#### Multi-scale grouping and models

The 11 nodes were grouped into four functional super-regions: Visual {V1, V2, V4}, Frontal {DLPFC, FEF}, Parietal {LIP, ACC, PCC}, and PM {P1, P2, P3} or {Y1, Y2, Y3}. The cross-scale consistency loss was applied to these multi-region groups. Two models were tested (i) MLP with weighted SNR loss and (ii) MLP with multi-resolution loss.

#### EC distribution and hub identification

Among two monkeys, the NPI-inferred EC showed a strong excitatory imbalance (Fig. 3a; 95.8% excitatory vs. 4.2% inhibitory), confirming that PM-to-cortex connectivity is predominantly excitatory. From NPI-inferred EC, we further identified the hub based on the combined in-degree and out-degree measures (Appendix C). Results from two monkeys suggested that FEF and DLPFC are two major hubs in this specific experiment, representing two key nodes in the visual and frontal areas, respectively.

**Figure 3:**
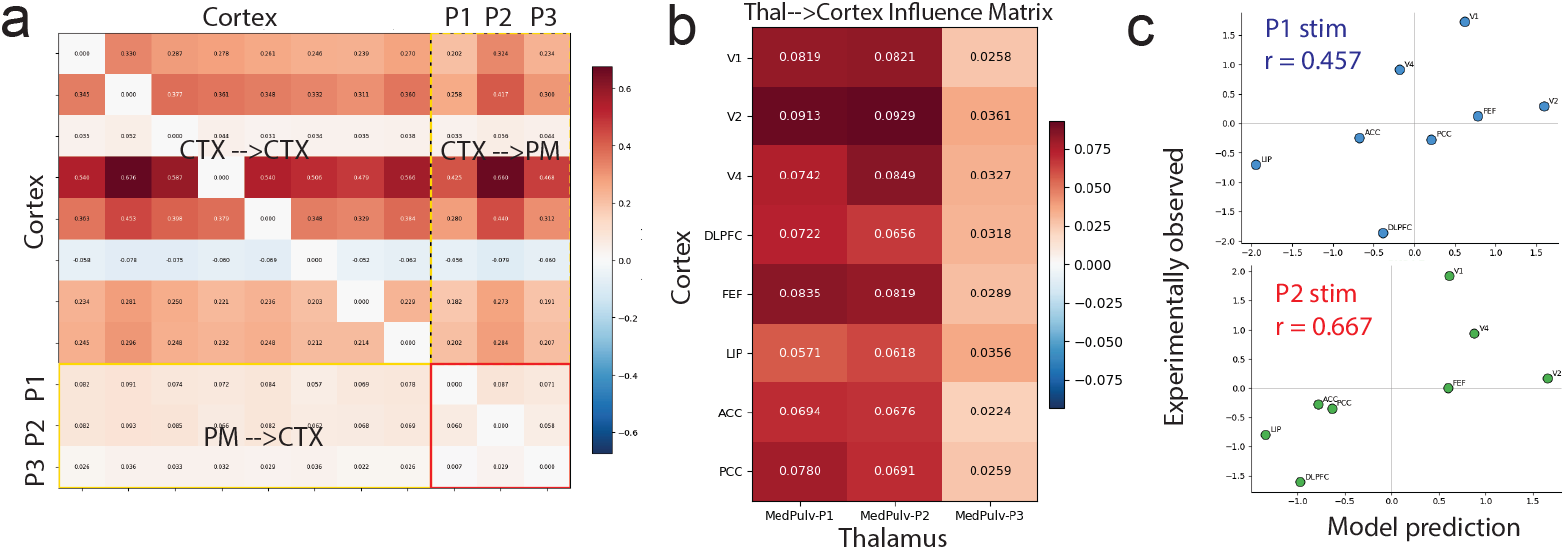
MacStim fMRI data results. (**a**) Dataset 1: Inferred EC matrix and connectivity graph (Monkey P). (**b**) Directed thal → cortex connectivity as an influence matrix. (**c**) Model-predicted vs experimentally-perturbed cortical responses (Monkey P), both P1 and P2 site-stimulations showed strong positive correlation. Results were obtained from the multi-resolution loss function.

#### Prediction of cortical responses

We further virtually perturbed the thalamic sites in the NPI-inferred thalamus-to-cortex influence matrix (Fig. 3b) and correlated the predicted output with the averaged change in cortical response induced by site-specific (P1/P2/P3 or Y1/Y2/Y3) thalamic stimulations. Results from Monkey P showed statistically significant correlation (Fig. 3c). More result figures from the second monkey are shown in Appendix C.

### 5.3 Dataset 2: Human Thalamocortical Resting-state fMRI Recordings

#### Data

The Human Thalamocortical fMRI dataset (HumanTC) consists of resting-state fMRI recordings of 12 human participants (eyes open). Subject IDs: sub-chon, sub-jche, sub-jmei, sub-ksei, sub-lsut, sub-lwan, sub-mdeb, sub-msco, sub-rmru, sub-sget, sub-tkas, sub-wkoo. Brain coverage: a total of 802 nodes: 360 cortical ROIs (HCP MMP 1.0 atlas; [25, 26]) plus 442 thalamic voxels (THOMAS parcellation; [16]), all in MNI152NLin2009cAsym space. Each subject provides 666 temporal samples in the resting period; data preprocessing included detrending and spatial smoothing (*σ*= 1), followed by z-scoring per region. Thalamic THOMAS groups: 21 named nuclei (AV, VA, VLa, VLp, VPL, Pul, LGN, MGN, CM, MD, Hb [lh only]; 297 of 442 voxels assigned; 145 boundary voxels excluded). Experimental procedure was approved by the Institutional Review Board (IRB) of Princeton University [27].

#### Model training and FC fingerprint reference

Four model configurations were considered for each subject: (i) MLP with standard loss, (ii) Dropout MLP with standard loss, (iii) DropoutMLP with weighted SNR loss; and (iv) DropoutMLP with multi-resolution loss. Because no direct thalamic stimulation was available, we considered an independent group-level thalamocortical FC reference (`thalamocortical_fpFC`) [28], which provides 180 left-hemisphere cortical ROIs × 442 thalamic voxels (Fig. 4a) [29]. Note that this reference was not used during model training (i.e., no leakage).

**Figure 4:**
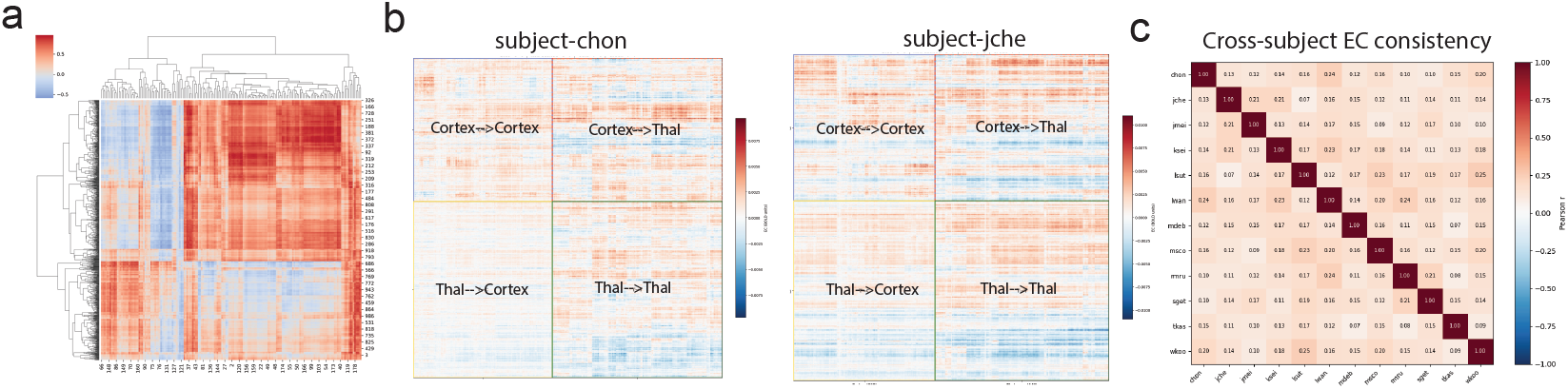
HumanTC fMRI data results. (**a**) Group-level thalamocortical FC reference (unpublished). (**b**) NPI-inferred 802 × 802 EC maps from two subjects. (c) EC map consistency across 12 subjects. Results were obtained from the weighted SNR loss function.

#### Assessment

For each subject, the empirical FC was computed as the Pearson’s correlation matrix derived from the resting-state fMRI signals across all 802 regions. The model-predicted FC was generated by iteratively running the trained model forward and computing the correlation structure of generated time series accordingly. Pearson’s *r* between the flattened empirical and model FC was then used as a reconstruction fidelity metric. As shown in Table 2 (2nd column), the DroputMLP outperformed standard MLP in all aspects, with the weighted SNR loss yielding the best reconstruction quality. Results of subject-wise FC quality and EC heatmap are shown in Appendix C.

**Table 2:** FC reconstruction and FC correlation comparisons based on different methods at the voxel level (*N* = 802). Mean and SD was computed across 12 subjects.

| Model | $r$ (FC reconstruction) | $r$ (w/ fingerprint FC) |
| --- | --- | --- |
| MLP, standard loss | $0.635 \pm 0.217$ | $0.205 \pm 0.128$ |
| DropoutMLP, standard loss | $0.741 \pm 0.077$ | $0.206 \pm 0.100$ |
| DropoutMLP, weighted SNR loss | $0.776 \pm 0.065$ | $0.285 \pm 0.090$ |
| DropoutMLP, Multi-resolution loss ($\alpha = 0.1$) | $0.743 \pm 0.047$ | $0.275 \pm 0.090$ |

We further compared the model FC with the existing thalamocortical FC (442 thalamic voxels ×180 ROIs), averaged across 4 resting-state datasets [29]. We extracted the thal → cortex block of each subject’s model FC and compared it with this reference. This provides an independent validation that does not use the subject’s own data. The weighted SNR loss again achieved the best alignment (Table 2, 3rd column), confirming that the NPI inferred biologically meaningful thalamocortical connectivity.

We also adopted THOMAS parcellation (*K* = 21) [16] and used Yeo-7 network parcellation [30] for the cortex (dividing the cortex into 7 functionally clusters). In this case, the thalamus is then defined at the nucleus level and the cortex is defined at the group level. We found that 19 of 21 THOMAS nuclei aligned positively with the FC reference (Table 3 in Appendix).

**Table 3:**
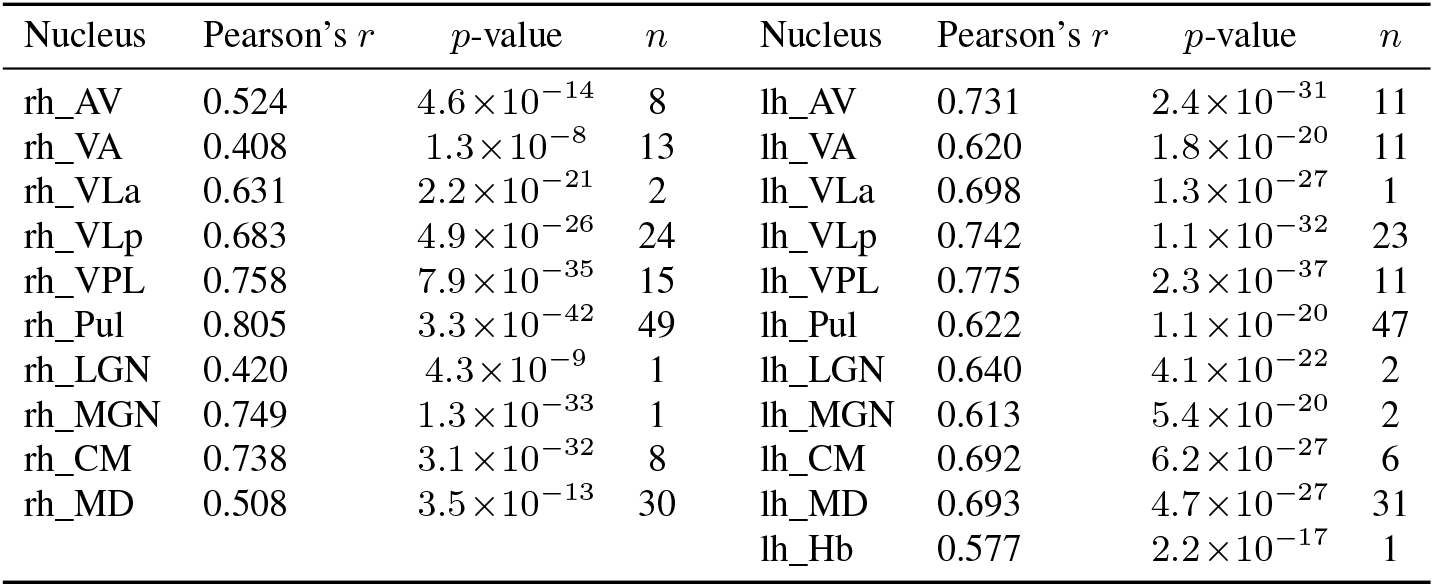
Thalamic nucleus FC reference correlation (*n* denotes the number of voxels within the thalamic nucleus).

Individuals’ inferred EC showed similar structures (see EC maps from two representative subjects in Fig. 4b). Due to the lack of EC ground truth, we assessed the consistency of NPI-inferred thalamocortical EC across 12 subjects (Fig. 4c). While there was shared EC structures between subjects in the off-diagonal, each subject’s EC matrix was also unique to identify the individual from the group (i.e., “personalized fingerprinting”).

Finally, at the voxel level, population-averaged thalamocortical EC showed a 63% excitatory and 37% inhibitory split in both directions (thalamus→ cortex and cortex→ thalamus), consistent with the known predominance of excitatory projections.

## 6 DBS Optimization via Thalamocortical EC

The NPI-inferred thalamocortical EC matrix 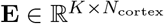 (*K* = 21 thalamic nuclei, *N*_cortex_ = 360 ROIs) encodes the predicted causal influence of each thalamic nucleus on every cortical region. Let **A** = **E**^*T*^ denote a thal → cortex *influence matrix*; column *k* is the cortical footprint of nucleus *k*. Under the assumption of a feedforward linear response model, the predicted cortical response to thalamic stimulation **u** ∈ ℝ^*K*^ (assuming as a DC constant) is **ŷ** = **Au**, which excludes the indirect effect through cortico-cortical EC. Given a desired change in the cortical target **y**^*∗*^ = **y**_post-stim_ −**y**_pre-stim_ (where the relative change in post-stimulation is computed by averaging the BOLD response across time and trials), we formulated the *sparse DBS target-selection problem*:

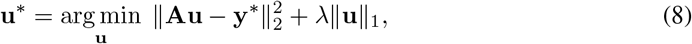

where the *l*_1_ penalty (as in LASSO [31]) favors sparse activation of thalamic nuclei, matching the clinical constraint of minimal implant sites.

For single or multiple thalamic site selection, we chose the optimal thalamic nuclei by the largest correlation of matching cortical responses: *k*^*∗*^ = arg max_*k*_ *ρ*(**A**_:,*k*_, **y**^*∗*^), with magnitude 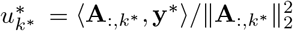. This reduces target selection to a nearest-neighbor search in nucleus footprint space. Because EC is inferred for each individual, the subject’s influence matrix **A**_*s*_ yields a different nucleus ranking. In contrast, a group-averaged EC would miss individual variability. In addition, the surrogate ANN model also enables nonlinear dosage simulation beyond the linear EC approximation; we can perturb the initial state **x**_0_ ← **x**_0_ + Δ_*k*_**e**_*k*_ and roll out 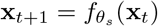 iteratively for *T* steps. As shown in Fig. 3c, the time-averaged cortical response approximately validates the linear response prediction, enabling multi-nucleus perturbation *in silico*.

We tested this linear control strategy using the setting from the MacStim dataset. The question is framed as: what is the optimal stimulation combinations (i.e., **u** = [*u*_1_, *u*_2_, *u*_3_], with the first/second/third element representing the stimulation amplitude for the P1/P2/P3 site in Monkey P’s PM area, respectively) to modulate two randomly selected cortical targets (e.g., both excited, or one excited and the other suppressed)? In experiments, we varied the cortical target locations and objectives and derived the approximate solutions. Our results showed comparable patterns (Fig. 5); but this strategy failed to turn off the off-target activity because of the cortico-cortical excitability.

**Figure 5:**
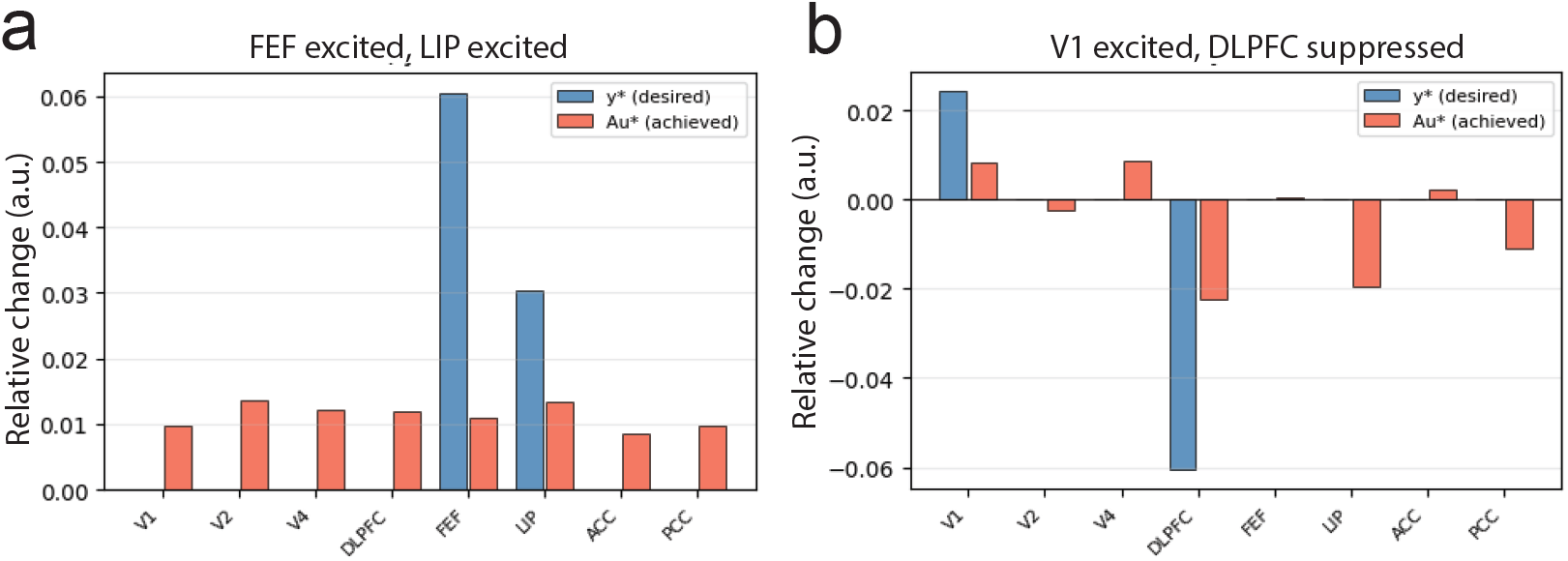
Comparison of desired target response y^*∗*^ and the controller response Au. (**a**) Dataset 2: both FEF and LIP excited. Subptimal control solution: **u**^*∗*^ = [0, 0, 0.375]. (**b**) V1 excited and DLPFC suppressed. Suboptimal control solution: **u**^*∗*^ = [−1.125, 1.823, −1.909].

## 7 Discussion

### SNR weighting and multi-resolution training

Our two proposed loss functions showed some advantages in EC estimation for synthetic and real fMRI data, especially where the thalamic nuclei have much lower SNR than the cortical regions. Theorem 1 provides formal justification. The weighted SNR loss is grounded by the heteroscedastic noise model (Propositions 1–2), showing improved generalization in real fMRI data. Multi-resolution training imposes the cross-scale consistency at the cost of increased complexity, and is likely to perform better when training samples are sufficient.

### From EC estimation to DBS optimization

The central premise of this work is that an accurate, subject-specific thalamocortical EC map is the key missing ingredient for data-driven DBS strategies. Our preliminary results provide a proof-of-concept and justification using causal thalamic perturbation data. The MacStim dataset provides the first NPI validation against *in vivo* perturbation responses. Some promising data were observed in two monkeys, but there was also some inconsistencies between monkeys and thalamic sites. More follow-up investigations and future experiments are still required.

The inferred EC provides biological constraints for personalized DBS. Once the EC is estimated, the choice of optimal stimulation target and magnitude reduces to a convex optimization problem with a well-posed solution. The linear control model and the nonlinear surrogate-based forward simulation provide complementary tools: the former for fast, interpretable thalamic target identification; the latter for more precise dosage planning. Our data-driven personalized neurostimulation framework is conceptually similar to the virtual brain twins motivated from a biophysical model [22]. Data-driven and biophysical models are complementary and may benefit from a hybrid model integration.

### Thalamocortical EC

In the MacStim dataset, we observed a predominant excitatory EC map, consistent with the known dominance of of glutamatergic thalamocortical synapses. In the HumanTC dataset, a population-averaged 63%/37% excitatory/inhibitory split was observed in the EC map. Note that the direct synaptic connections from the pulvinar to the cortex are traditionally considered to be excitatory, FC between them can be functionally inhibitory (acting as a gate for cortical information). While the exact EC ground truth of the large thalamocortical network remains unknown, our results showed some consistent observations at both voxel level and nucleus level (THOMAS parcellation), and the CM nucleus showed the strongest individual-nucleus correspondence (*r* ≈ 0.83 bilaterally).

## 8 Conclusion

We present a surrogate brain framework for estimating thalamocortical EC and use it for personalized thalamic DBS optimization. We proposed DropoutMLP and two new loss functions to address the statistical challenges of high-dimensional thalamocortical surrogate training given heterogeneous SNRs in resting-state fMRI signals. Our approach identified individualized thalamocortical EC map from resting-state fMRI datasets from 12 humans and 2 macaque monkeys. The personalized thalamocortical EC directly leads to a sparse thalamic DBS target-selection strategy for distributed cortical modulation. While the surrogate ANN further enables nonlinear in-silico stimulation, the convex optimization solution yields a fast, computationally efficient parameter selection procedure for linear control used in clinical applications. Together, our work here represents a proof-of-concept for precise thalamocortical circuit mapping and personalized thalamic DBS stimulation.

## 9 Limitations

In the MacStim dataset, only the medial pulvinar site was perturbed via infrared stimulation, yet a full reconstruction of thalamocortical EC will benefit from stimulations of cortical sites. The HumanTC dataset has small numbers of voxel counts in thalamic nuceli (e.g., rh_LGN (1 voxel), rh_VLa (2), lh_VLa (1), and lh_Hb (1)), and the THOMAS atlas does not subdivide the pulvinar into functional subregions, all contributing to unreliable single-subject thalamocortical EC estimates.

We have assumed the linear neural response in thalamic DBS control. However, in neurostimulation, the relationship between stimulation amplitude and the resulting neural response is not linear. While low-amplitude stimulation typically causes direct, low-latency excitation of nearby neurons, increasing the amplitude often shifts the balance toward inhibition. The current thalamocortical model has not considered thalamic inhibition from the thalamic reticular nucleus (TRN) and has ignored cortico-cortical feedback. Additionally, DBS operates at a millisecond timescale that cannot be directly captured by slow fMRI-BOLD signals.

Unlike structural connectivity that is relatively stable, it remains unclear how resting-state EC may change in task conditions where the brain state is influenced by both bottom-up and top-down contributions. Finally, mapping from fMRI-inferred EC to electrophysiological stimulation effects requires empirical calibration against concurrent electrophysiology and neurostimulation data. Future work can use this framework to study stimulation-induced thalamocortical plasticity during learning. Appendix D briefly discusses the broader impact of our work.

## Acknowledgments

We thank Michael Arcaro and Xingyu Liu for sharing the human fMRI data and feedback. This work was partly supported by the National Institute of Health grants R01-MH139352, P50-MH132642 and RF1-DA056394 to Z.S.C.

## A Detailed Proofs

### A.1 Proof of Proposition 1

The SNR-weighted estimator minimizes 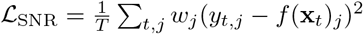. Taking the derivative with respect to *f* (**x**)_*j*_ and setting it to zero gives the normal equations of a weighted least squares (WLS) problem. Since *E*[*y*_*t,j*_] = *f*^*∗*^(**x**_*t*_)_*j*_ (from the noise model in Eq. 5), the estimator is unbiased. The unbiasedness follows from the fact that *w*_*j*_ does not depend on the residuals. The variance calculation follows from the standard WLS formula: 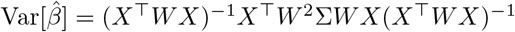, where 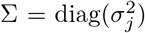 and *W* = diag(*w*_*j*_). Under the model 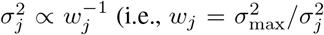, the optimal weighting), this reduces to (*X*^*T*^*WX*)^*−*1^, which is smaller than the unweighted (*X*^*T*^*X*)^*−*1^ in the positive-semidefinite order. Our choice *w*_*j*_ = (1−*λ*) + *λh*(tSNR_*j*_*/* max_*k*_ tSNR_*k*_) is a monotone approximation to the optimal weighting.

### A.2 Proof of Proposition 2

It follows from the standard Rademacher generalization bound [23] and applied to the weighted loss class *l*_*j*_(*f*, **x**, *y*) = *w*_*j*_(*f* (**x**)_*j*_ − *y*_*j*_)^2^. The Lipschitz constant of the squared loss is 2 · max_*j*_(*w*_*j*_) · max_*f*,**x**_ |*f* (**x**)_*j*_ − *y*_*j*_|, and the standard contraction lemma gives the stated bound.

A corollary naturally follows when combining SNR weighting and nucleus constraints. Specifically, let ℱ have output-layer trace-norm bound ∥*W*_out_ ∥_*∗*_ ≤ *B* and let *K* nucleus-level consistency constraints and SNR weighting with minimum weight *w*_min_ = 1− *λ* be applied. Then the excess risk satisfies:

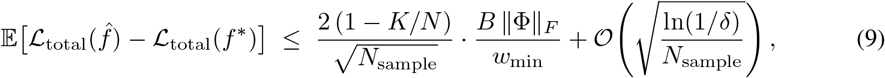

where Φ is the feature matrix. Smaller *K/N* (i.e., fewer constraints) or larger *w*_min_ (i.e., stronger SNR weighting) reduces the bound.

### A.3 Full Proof of Theorem 1

Let *F* be the hypothesis class of fine-model functions *f*_*θ*_ : ℝ^*N*^ → ℝ^*N*^ (e.g., *N* = 802 in Dataset 2), with output decomposed as 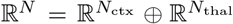 (e.g., *N*_ctx_ = 360, *N*_thal_ = 442).. Multi-resolution training augments the fine model with a coarse surrogate 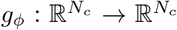(*N*_*c*_ = 381 after THOMAS grouping) and a cross-scale consistency loss:

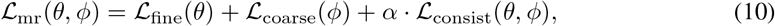

where *α* = 0.1 and 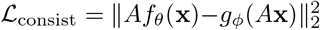 with 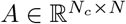 as the parcellation aggregation matrix. Additionally, the consistency term restricts to the sub-class

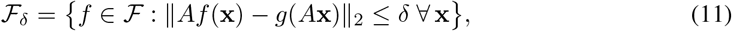

so *F*_*δ*_ ⊆ *F* for any *δ >* 0.

We elaborate on the uniform distribution assumption. Apply singular value decomposition (SVD) to *W*_out_ = *U* Σ*V* ^*T*^, where Σ = diag(*σ*_1_, …, *σ*_*d*_) and ∥*W*_out_∥_*∗*_ = Σ_*k*_ *σ*_*k*_ ≤ *B*. The uniform assumption states *σ*_*k*_ = *B/d* for all *k*. Each nucleus constraint ⟨**a**_*k*_, *W*_out_***ϕ***⟩ = *c*_*k*_ is equivalent to pinning the projection 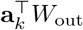, which removes a budget of *B/d* from the trace norm. After *K* constraints, the budget is (1 − *K/d*)*B* ≤ (1 − *K/N*)*B* because of *d* ≤ *N*.

By the standard trace-norm Rademacher bound, for a class of functions of the form *f* (**x**) = *W*_out_*ϕ*(**x**) with ∥*W*_out_∥_*∗*_ ≤ *B*:

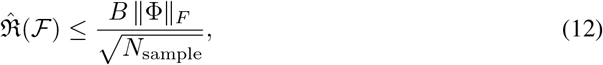

where 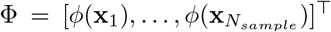. Each of the *K* consistency constraints pins one linear combination of the rows of *W*_out_ to a fixed value determined by the coarse model. Pinned directions contribute zero variance to the Rademacher supremum:

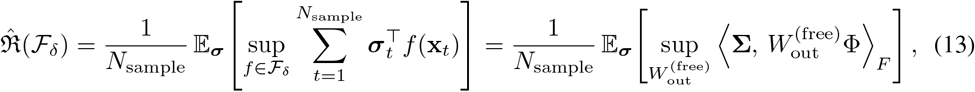

where 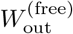 ranges over the (*N*− *K*) unconstrained output rows, and **Σ** is the Rademacher matrix restricted to those rows. Under the uniform trace-norm assumption, the budget available to the free rows is (1 − *K/N*) *B*. Substituting into (12) gives (7). Since the standard generalization bound gives 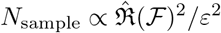, it follows that *N*_multi*−*resolution_ ≤ (1 − *K/N*)^2^ *N*_sample_.

## B Training Details

All models were trained exclusively on simulated or resting-state fMRI-BOLD signals. Assessment were evaluated on held-out samples. In every setting the training objective is one-step-ahead prediction in the NPI procedure.

### B.1 Synthetic Data (RNN simulation)

#### Network

The synthetic thalamocortical RNN comprises *N*_thal_ = 15 thalamic voxels and *N*_cort_ = 35 cortical ROIs (*N* = 50 total). Each scenario generated 8,000 time-steps; the sliding-window pipeline (stride *s* = 3) yields 7,997 input–output pairs, split 80/20 into 6,397 training and 1,600 test samples in 5-fold cross-validation.

#### Network architecture

The ANN was an MLP with three fully connected layers and ReLU activation functions:

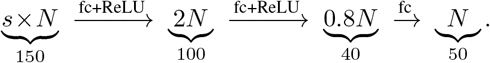

where “fc” denotes fully connected nodes. The paired coarse model used the same formula with *N*_merged_ = 43 (seven thalamic adjacent-voxel pairs + one thalamic singleton + 35 cortical ROIs).

#### Optimization

Adam optimizer; learning rate lr = 2.5 × 10^*−*4^, weight decay regularization *l*_2_ = 5×10^*−*5^; batch size 64. Default perturbation strength Δ = 1.0 (z-scored signals have unit variance), but the Δ value was also tried on 0.01 and 0.1 to test the model robustness.

#### SNR scenarios

Thalamic tSNR was drawn uniformly from [3, 8] (moderate) or [2, 5] (severe); cortical tSNR was drawn uniformly from [15, 30]. In the uniform baseline condition, we used a fixed tSNR = 15 for all nodes. In SNR-weighted training, we used a log-normalized weighting strategy (*w*_*j*_ = (1 − *λ*) + *λ* · log(1 + tSNR_*j*_*/*tSNR_max_)*/* log 2, *λ* = 0.3).

### B.2 Dataset 1 (MacStim)

#### Preprocessing

Raw fMRI-BOLD signals (amplitude range ∼ 1470–2090) were mean-demeaned independently per region prior to training: the temporal mean of each region was subtracted, with no division by standard deviation. The per-region standard deviation was retained separately to compute tSNR and perturbation strength. Region-level tSNR (defined as mean/std on raw BOLD before demeaning): PM = 68.23 (lowest); cortical ROIs 107–144. Perturbation strength was set to 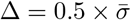, where 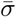 denotes the mean standard deviation averaged across all 11 regions. The same three training variants (standard, SNR-weighted, and multi-resolution) were applied independently and identically to both Monkey P and Monkey Y using the same hyperparameters.

#### Network architecture

The ANN was a three-layer MLP with Tanh activation functions and *N* = 11 nodes: 8 cortical ROIs (V1, V2, V4, DLPFC, FEF, LIP, ACC, PCC) and 3 thalamic PM sites treated as independent nodes (indexed 8, 9, 10):

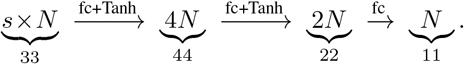

This yields ≈ 2,739 trainable parameters. Given 237 input–output pairs derived from 240 resting-state time points (sliding window, *s* = 3) with an 80%/20% train–test split, 189 training samples were available, yielding a parameter-to-sample ratio of ∼14.5 : 1.

#### Optimization

Adam optimizer; learning rate lr = 2.5 × 10^*−*4^, weight-decay regularization *l*_2_ = 5 × 10^*−*5^; batch size 32; 200 epochs; train/test split 80%/20%.

#### Coarse grouping

The 11 nodes were partitioned into **three** functional super-regions for the cross-scale consistency loss: Visual {V1, V2, V4}, Frontal+Parietal {DLPFC, FEF, LIP, ACC, PCC}, and Thalamic {PM-site 1, PM-site 2, PM-site 3}, yielding *N*_*c*_ = 3 coarse nodes. The coarse companion model shared the same Tanh-MLP structure with output dimension *N*_*c*_ = 3:

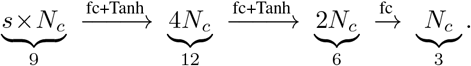

The cross-scale consistency loss (*α* = 0.1) was applied between the fine model’s group-averaged predictions and the coarse model’s outputs.

#### Hub identification

Given the inferred 11×11 EC matrix, nodal centrality was computed as the sum of out-degree (row-sum of |EC |) and in-degree (column-sum of| EC|). The two cortical nodes with the highest combined degree were designated as network hubs.

| Region (Monkey) | Out-deg | In-deg | Total |
| --- | --- | --- | --- |
| FEF (P) | 3.6861 | 1.9272 | 5.6133 |
| DLPFC (P) | 5.4471 | 1.7444 | 7.1915 |
| V1 (P) | 2.6705 | 2.0097 | 4.6802 |
| FEF (Y) | 3.7842 | 2.9614 | 6.7456 |
| DLPFC (Y) | 2.5377 | 2.5115 | 5.0492 |
| V1 (Y) | 1.1576 | 1.8010 | 2.9586 |

### B.3 Dataset 2 (HumanTC)

#### Subjects

Twelve human participants were included: sub-chon, sub-jche, sub-jmei, sub-ksei, sub-lsut, sub-lwan, sub-mdeb, sub-msco, sub-rmru, sub-sget, sub-tkas, sub-wkoo. One surrogate model was trained independently for each subject.

#### Signal and preprocessing

Resting-state BOLD was acquired for each subject across *T* = 666 time points. Cortical activity was summarized using the Human Connectome Project Multi-Modal Parcellation (HCP-MMP1.0) atlas, yielding *N*_cort_ = 360 parcel-averaged time-series. Thalamic activity was represented at full voxel resolution using a subject-space thalamic mask, yielding *N*_thal_ = 442 voxel time-series. Cortical signals were provided as a detrended, spatially smoothed (*σ*= 1) parcel-mean CSV; thalamic signals as a corresponding .npy array. The two modalities were concatenated to form the joint fine-grained signal **s**(*t*) ∈ ℝ^*N*^, *N* = 802.

#### THOMAS nucleus grouping

Thalamic voxels were mapped to named nuclei using the THOMAS atlas (`thal_THOMAS.nii.gz`). For each voxel in the thalamic mask, its MNI world coordinate was projected into THOMAS atlas space and the nearest integer label was retrieved. This yielded 21 named nucleus groups across both hemispheres: 10 right-hemisphere nuclei (AV, VA, VLa, VLp, VPL, Pul, LGN, MGN, CM, MD) and 11 left-hemisphere nuclei (AV, VA, VLa, VLp, VPL, Pul, LGN, MGN, CM, MD, Hb). A total of 297 of the 442 thalamic voxels were assigned to these named groups; the remaining 145 boundary voxels received no label and were excluded from the coarse-scale training signal and loss. Nucleus sizes ranged from 1 to 49 voxels (median = 11). The coarse thalamic signal for each nucleus was the mean time-series across its member voxels; cortical parcels remained at singleton granularity. This produced a coarse-resolution signal 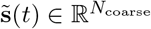 With *N*_coarse_ = 381 (360 cortical ROIs + 21 nucleus groups). Additionally, we adopted Yeo-7 network parcellation [30] that divides the cortical ROIs into 7 functionally clusters, yielding a compact (28 × 28) thalamocortical network representation.

#### Network architecture and four model variants

All surrogate models share a three-hidden-layer multi-layer perceptron (MLP) architecture. The input is a temporal stack of *s* = 3 consecutive frames, so the effective input dimension is *s* × *N*. Four variants were trained per subject:

1. **Standard NPI.** Baseline model using the original NPI architecture (`ANN_MLP`): no dropout, no cross-scale supervision. Input → hidden 128 → hidden 64 → output *N*.
2. **DropoutMLP.** Same depth but with Bernoulli dropout (*p* = 0.3) after each hidden ReLU activation; dropout is active only during training and disabled during inference (`model.eval()`).

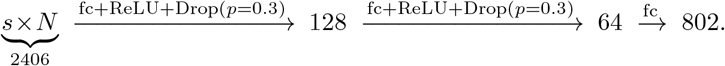

Combining dropout with *l*_2_ weight-decay regularization aimed to mitigate overfitting.
3. **SNR-Weighted.** Same `DropoutMLP` architecture but with per-region loss weighting during training. The SNR proxy for each of the 802 regions was defined as the inverse of its temporal standard deviation:

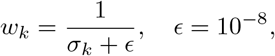

where *σ*_*k*_ = std_*t*_[*s*_*k*_(*t*)]. Weights were log-scaled with strength 0.3 before being applied to the per-region squared errors, down-weighting high-variance (low-SNR) thalamic voxels relative to cortical parcels (mean cortical SNR proxy 1.32, mean thalamic SNR proxy 1.0).
4. **Multi-Resolution (MR).** A coupled training scheme involving two simultaneously trained models: the fine-grained `DropoutMLP` (*N* = 802) and a coarse `DropoutMLP` (*N*_coarse_ = 381) with identical depth and hidden dimensions. The fine model received an auxiliary cross-scale regularization term: for each of the 17 THOMAS nucleus groups containing more than one voxel, the voxel-mean of the fine model’s thalamic predictions was compared to the voxel-mean of the targets at the same spatial scale. Cortical singletons and the four single-voxel nucleus groups (rh-LGN, rh-MGN, lh-VLa, lh-Hb) did not contribute to this term. The total fine-model loss was:

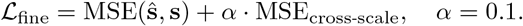

The coarse model was trained independently on the nucleus-averaged signal with standard MSE.

#### Data preparation

For each subject, the joint signal was converted to a multi-step supervised learning problem using a lag of *s* = 3 frames: each input is [**s**(*t*− 2), **s**(*t* −1), **s**(*t*)] and the target is **s**(*t* + 1), yielding 663 input-output pairs per subject. An 80/20 chronological data split was applied, yielding 529 training samples and 134 test samples.

#### Optimization

All four variants used the Adam optimizer with learning rate *η* = 3 ×10^*−*4^ and *l*_2_ weight decay regularization parameter *λ* = 1 × 10^*−*3^, batch size 64, and a maximum of 500 epochs. Early stopping monitored the held-out validation loss (patience = 40 epochs, *δ* = 10^*−*4^); the checkpoint with lowest validation loss was restored at the end of training..

#### EC estimation

After training, NPI virtual perturbations were applied to each fine model to estimate EC. Each of the 802 nodes was perturbed individually; the perturbation magnitude was set to

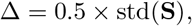

where std(**S**) denotes the global standard deviation of the entire *T* × *N* normalized BOLD matrix, computed independently for each subject (Δ ranged from 0.44 to 0.47 BOLD units across subjects). Self-connections were zeroed post-hoc. This produced a 802 × 802 directed EC matrix capturing Thalamus →Cortex, Cortex →Thalamus, Cortex →Cortex, and Thalamus →Thalamus pathways at full voxel resolution. All analyses retained the 442 thalamic voxels individually rather than collapsing to nucleus representatives, preserving fine-grained spatial organization within the thalamus.

## C Additional Results and Figures

In the synthetic dataset, Figure 6 presents additional results on RMSE, model robustness with respect to the perturbation strength and the noise level.

**Figure 6:**
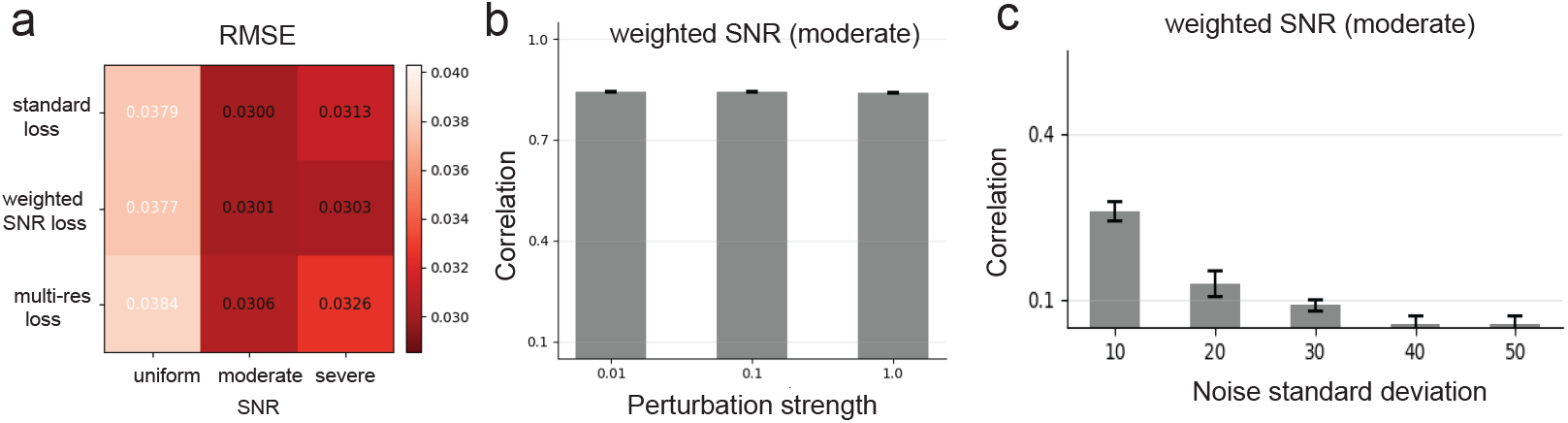
Synthetic data results. (**a**) Mean RMSE comparisons under three SNR conditions. (**b**) Estimation was robust with perturbation strength Δ. (**c**) Correlation degraded with increasing level of noise. All simulations were done for a 50-node network.

In the MacStim dataset, Figure 7 shows representative learning curves of three different loss functions. Additionally, Figure 8 shows the model prediction results of Monkey Y (compared to results of Monkey P in Fig. 3).

**Figure 7:**
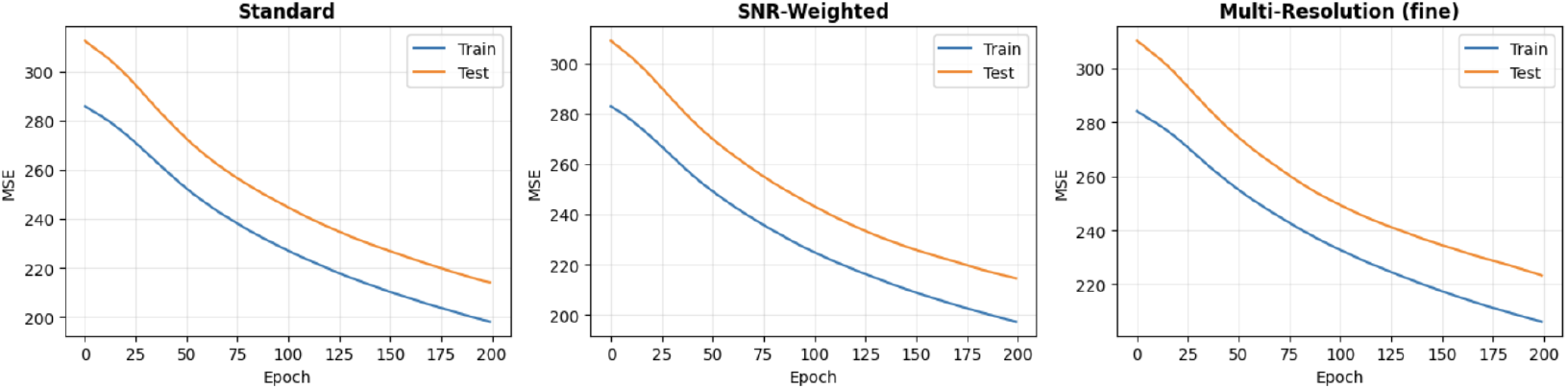
MacStim Dataset results. MLP learning curves for training and testing samples over 200 epochs for the standard, SNR-weighted, and multi-resolution loss functions.

**Figure 8:**
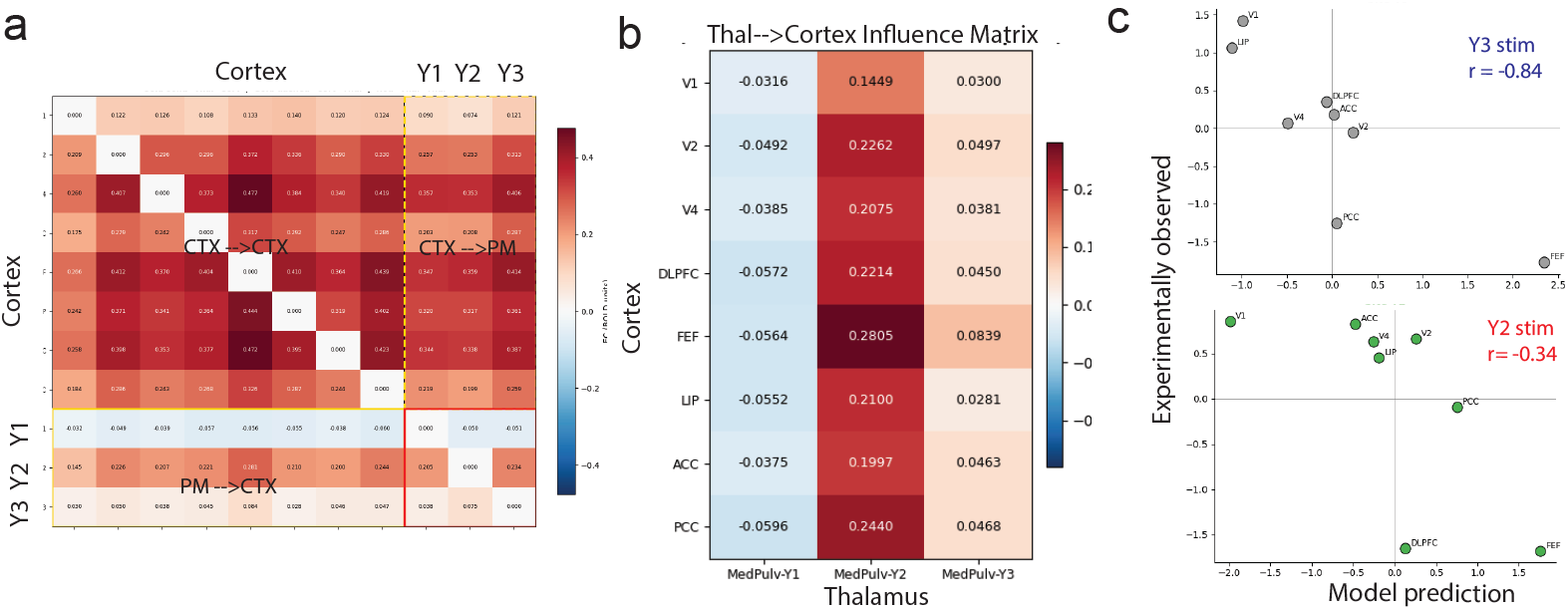
MacStim Dataset results for Monkey Y. (**a**) Inferred EC matrix and connectivity graph (Monkey P). (**b**) thal→cortex connectivity as an influence matrix. (**c**) Model-predicted vs experimentally-perturbed cortical responses. Interestingly, the model prediction still showed significant correlation with the experimental observations, but with a negative correlation (compared to Monkey P), suggesting a highly subject-specific DBS effect.

In the HumanTC dataset, Figure 9 presents the FC reconstruction fidelity comparison between four tested models across 12 subjects. Figure 10 presents the visualization of a 28 × 28 thalamocortical EC map based on THOMAS parcellation and Yeo-7 network. Table 3 summarizes the detailed results for thalamic nucleus FC reference correlation. Because of multiple comparisons, *p*-values were adjusted using Bonferroni correction.

**Figure 9:**
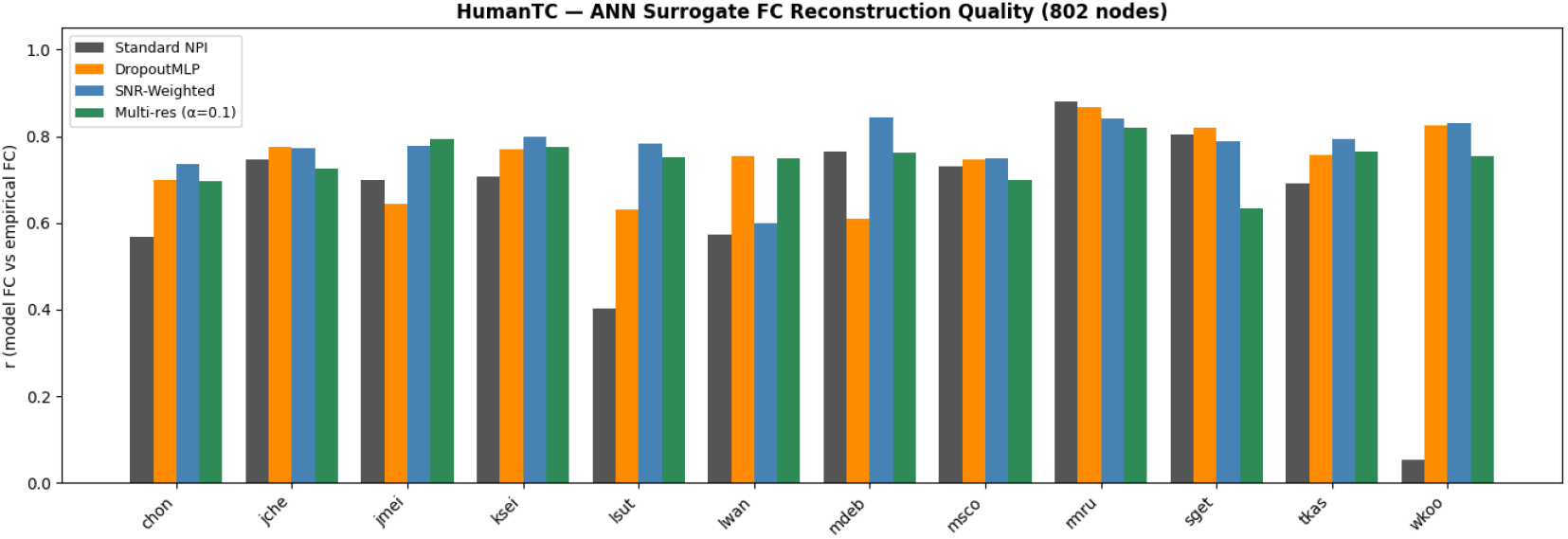
HumanTC Dataset results. NPI-inferred model FC reconstruction fidelity (vs. empirical FC) across 12 subjects. SNR-weighted loss function produced overall best result, whereas the standard NPI (standard loss) produced the the worst results, especially in subjects lsut and wkoo.

**Figure 10:**
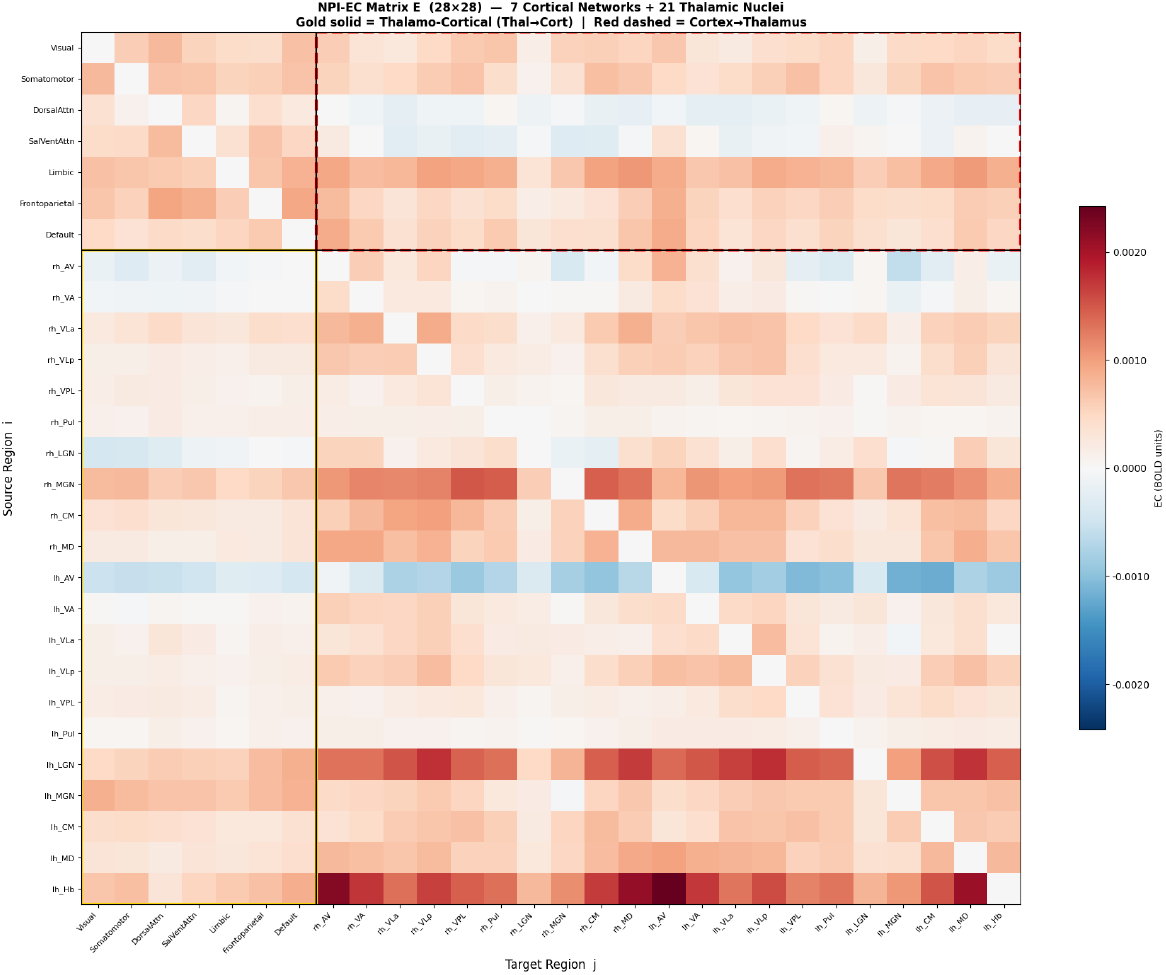
HumanTC Dataset results. Visualization of a compact 28×28 thalamocortical EC map based on THOMAS parcellation (21 nuclei) and Yeo-7 network (7 cortical functional clusters).

## D Broader Impact

This work contributes to the intersection of computational neuroscience, NeuroAI, and neurotechnology by introducing a surrogate-brain framework for estimating individualized thalamocortical effective connectivity and optimizing DBS strategies from resting-state fMRI data. Beyond advancing scalable causal modeling of large-scale brain networks, the proposed framework may support future development of personalized neuromodulation, digital twin brain, and adaptive closed-loop neurostimulation systems for neurological and psychiatric disorders. More broadly, this work illustrates how AI-based surrogate models can serve as mechanistic tools for probing and controlling distributed biological systems. Important limitations and ethical considerations remain, including uncertainty in fMRI-derived EC, potential dataset bias, inter-individual variability, privacy concerns surrounding personalized brain models, and the need for rigorous clinical validation before translational deployment. Our framework is intended as a proof-of-concept computational approach rather than a clinically validated DBS planning system, and future work will require integration with electrophysiology, neurostimulation experiments, and regulatory oversight to ensure safe and responsible clinical applications.

